# Smart Slides for Optical Monitoring of Cellular Processes

**DOI:** 10.1101/2023.08.03.549853

**Authors:** Julia Ackermann, Eline Reger, Sebastian Jung, Jennifer Mohr, Svenja Herbertz, Karsten Seidl, Sebastian Kruss

## Abstract

The molecules released by cells are a fingerprint of their current state. Methods that measure them with high spatial and temporal resolution would provide valuable insights into cell physiology and diseases. Here, we develop a nanosensor coating that transforms standard cell culture materials/dishes into “Smart Slides” capable of optically monitoring biochemical efflux from cells. For this purpose, we use single wall carbon nanotubes (SWCNTs) that are fluorescent in the beneficial near-infrared (NIR, 850 – 1700 nm) window. They are chemically tailored to detect the neurotransmitter dopamine by a change in fluorescence intensity. These nanosensors are spin-coated on glass substrates and we show that such sensor layers can be sterilized by UV light and can be stored in dry condition or buffer for at least 6 weeks. We also identify the optimal sensor density to maximize sensitivity. Finally, we use these materials to image dopamine release from neuronal cells cultivated on top in the presence of various psychotropic substances, which represents a system to test pharmaceuticals for neurological or neurodegenerative diseases. Therefore, Smart Slides are a powerful tool to monitor cellular processes in cell culture systems.

## 1. Introduction

Cells release molecules and the spatiotemporal concentration profile is a biochemical fingerprint of the biological state.^[1,2]^ Real-time monitoring of these molecules is therefore important to gain insights into cell physiology and how cells respond to changes in their environment such as nutrient availability, environmental toxins, or drug exposure.^[3,4]^

Degeneration and death of neurons leading to a decline of cognitive and motor functions, can be caused by a variety of factors, which is the reason why neurodegenerative disorders are currently incurable and represent a significant burden to society^[5,6]^. To understand such complex mechanisms and develop effective treatments, highly sensitive biosensors that detect temporarily- and spatially-resolved biomolecular changes at the cellular level are of great importance. For example, numerous methods have been developed to assess neurobiological processes by quantifying dopamine levels, which gives deeper insights into diseases such as Parkinson’s, depression, and addiction. Such methods include electrochemical and fluorescence-based methods or imaging mass spectrometry, all of which have their own advantages and disadvantages.^[7,8]^ However, to date a simple and ready-to-use tool for standardized *in vitro* cell culture to obtain dynamic information about the biological state of cells is missing.^[4]^

Biosensors based on nanomaterials have shown many advances in the past years.^[9,10]^ One example is single-wall carbon nanotubes (SWCNTs), which represent nanoscale building blocks with a versatile surface chemistry.^[11]^ Due to their fluorescence emission in the near infrared (NIR, 850 – 1700 nm) biological transparency window, imaging and sensing applications benefit from reduced scattering and autofluorescence of biological samples and thus provide a high signal-to-noise ratio.^[12,13]^ In addition, unlike many other fluorophores, they have excellent photostability and are suitable for long-term experiments.^[14]^ Upon chemical surface functionalization they are able to detect different biomolecules with high sensitivity even at the single-molecule level.^[15–17]^ For example, sensors have been developed for the detection of stress in plants^[18–20]^, proteins^[21,22]^, cancer markers^[23]^, bacteria^[24]^, or neurotransmitters^[25–27]^. Moreover, with stable SWCNT functionalization, these sensors exhibit very promising biocompatibility.^[28,29]^ Neurotransmitter detection has been demonstrated with a high spatial and temporal resolution for chemical mapping of cellular release processes of dopamine^[17,30–32]^ and serotonin.^[26]^ Recent developments also show that this technology can be tailored for common microscopy equipment^[33]^ and that extracellular dopamine can be reported in 3D by combining time-correlated single photon counting and confocal fluorescence microscopy.^[34]^ Moreover, SWCNT tracking can reveal the nanoscale extracellular space in brain tissue.^[35]^

However, to enable standardized biological experiments, it is crucial to integrate such sensors into common labware and microscopy settings. SWCNTs offer a distinct advantage in this regard, as they can form high-density sensor arrays. Stapelton *et al*. developed a SWCNT platform for studying nitric oxide by employing an avidin-biotin interaction as SWCNT immobilization strategy.^[36]^ This approach enabled them to immobilize higher concentrations of SWCNTs compared to the standard surface silanization with (3-aminopropyl)triethoxysilane (APTES), resulting in enhanced fluorescence signals. The main challenge is not to measure the fluorescence intensity itself but to ensure an even distribution of sensors on the sample. In general, prolonged physisorption times of sensors can lead to their aggregation, which complicates coating processes.

To address this issue, we present a method for converting glass substrates (glass bottom petri dishes) into “Smart Slides” by spin coating SWCNT sensors on surfaces to achieve a homogeneous sensor coating (**Figure 1 a**). The slides continue to support cell adhesion but also become “smart” by optically monitoring biochemical cell responses. We test the optimal conditions for analyte sensitivity and determine the functionality of the slides in terms of storage and sterilization conditions (**Figure 1 b**). In addition, we demonstrate the potential for drug testing by studying dopamine release in a neuronal cell model. Overall, we aim to establish SWCNT-based fluorescent sensor coatings as powerful material/tool for the life sciences.

**Figure 1.**
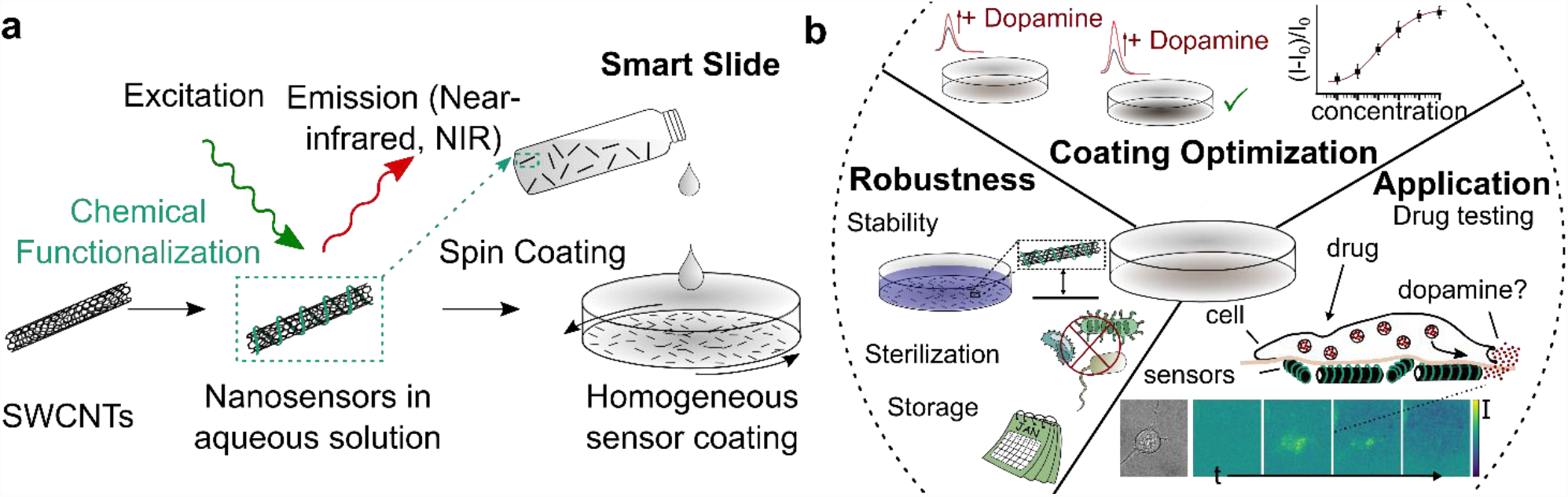
Smart Slides to image cellular responses by light. a) NIR fluorescent SWCNTs are rendered sensitive to the neurotransmitter dopamine by surface functionalization with specific ssDNA sequences. These nanosensors are coated on glass substrates (petri dishes) via spin coating. b) These sensor-coated substrates (Smart Slides) are optimized in terms of robustness for cell-based assays. They serve as versatile tools to assess the impact of pharmaceuticals on dopamine release by neurons.

## 2. Results and Discussion

### 2.1. Optimization of sensor coating and analyte sensitivity

Dopamine plays an important role in various neurodegenerative and mental health diseases. Therefore, we aimed to develop and optimize a dopamine-sensitive coating for cell culture studies. For this purpose, SWCNTs were functionalized with single-stranded (ss)DNA (GT)_10_ oligonucleotides, which non-covalently adsorb onto the SWCNT surface by π-π stacking interactions.^[37]^ Such (GT)_10_-SWCNTs are known to exhibit a fluorescence increase in response to dopamine.^[26,33,38]^ The prepared (GT)_10_-SWCNTs showed individual and narrow peaks in the absorption spectra, which indicates that they are well-dispersed (**Figure S1 a**). Their fluorescence also increased after addition of dopamine, confirming their sensitivity (**Figure 2 b**).

**Figure 2.**
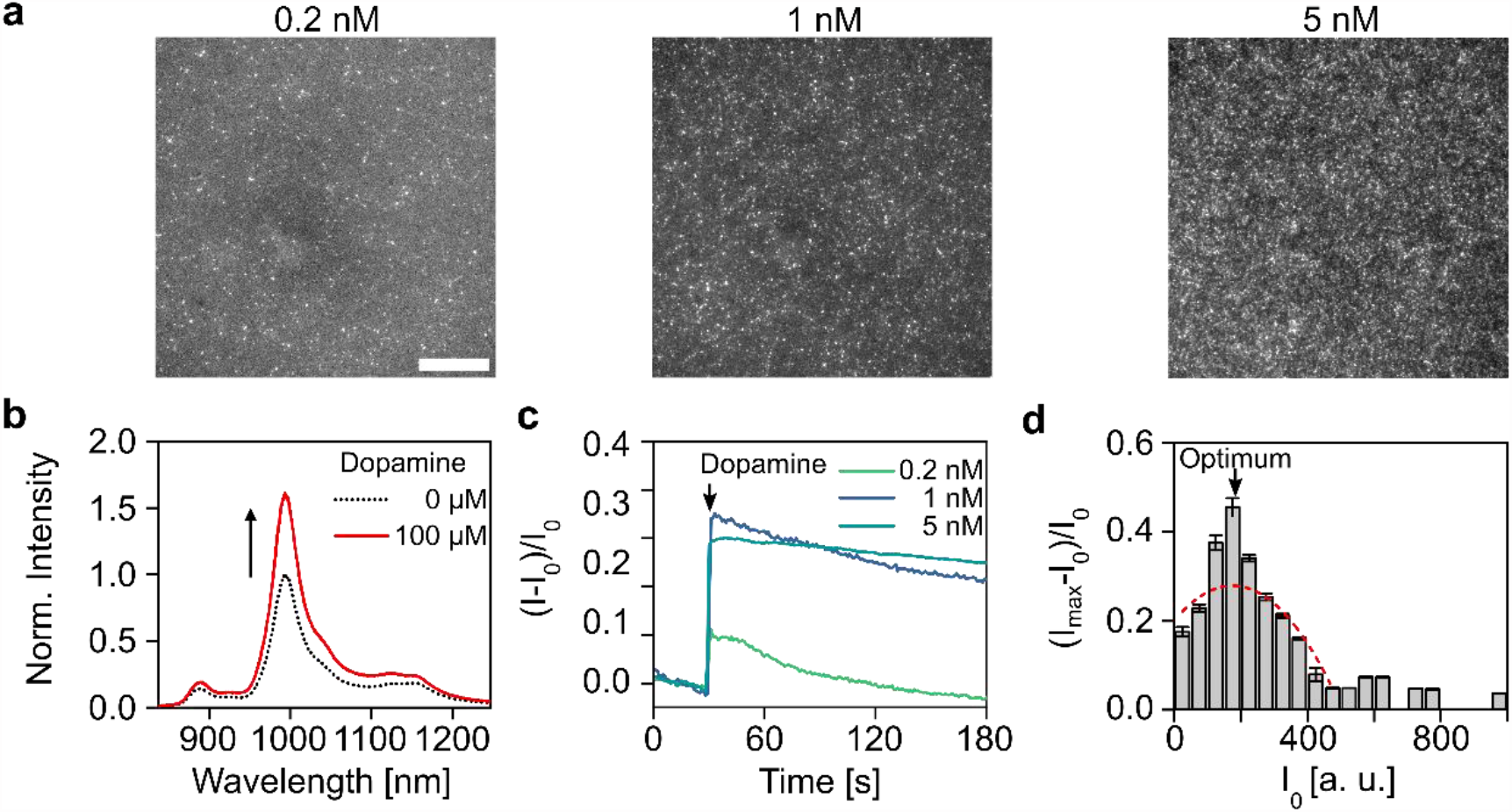
Optimal SWCNT density for Smart Slides. a) NIR fluorescence images of (GT)_10_-SWCNTs spin-coated at 1600 rpm for different SWCNT suspension concentrations. Scale bar = 20 μm. b) Normalized NIR fluorescence spectra of (GT)_10_-SWCNTs before and after addition of 100 μM dopamine in PBS. c) Real-time fluorescence changes of SWCNT coatings of different densities upon addition of 100 μM dopamine. d) Sensitivity of sensor coatings in response to 100 μM dopamine as a function of the initial SWCNT fluorescence intensity I_0_ of (GT)_10_-SWCNT coatings in PBS (mean ± SE, n ≥ 8). The red curve represents a model that accounts for density-dependent photophysical effects (see formula (1), R^2^ value of >0.95).

To achieve a homogeneous coating of SWCNTs on glass slides or glass-bottom petri dishes, spin coating was performed. For this purpose, the substrates were first modified with aminosilanes (APTES) to increase the electrostatic interactions between the SWCNTs and the glass surface. Subsequently, various parameters, such as SWCNT concentration (0.2 – 5 nM) and rotational speed (400 – 2000 rpm), were tested to optimize adsorption. The coating at the same SWCNT concentrations at different rotational speeds did not differ except for SWCNTs with a concentration of 5 nM (**Figure S2**). Consequently, we decided to use rotational speeds of 1600 rpm at which the distribution was homogeneous for all tested SWCNT concentrations and for which coating density could be adjusted by the SWCNT concentration (**Figure 2 a**). In addition, (GT)_10_-SWCNT coated surfaces exhibited a low defect ratio, as evidenced by the Raman G/D intensity ratio of 9.9 (**Figure S3**).^[39]^ Furthermore, we tested whether homogeneous coating is also possible for chirality-pure SWCNTs, which are more prone to aggregation.^[40]^ For this purpose, (GT)_10_-(6,4)-SWCNTs with a concentration of 0.1 nM (**Figure S1 b**) were spin-coated in a series of six consecutive coating cycles on the substrate. Again, a dense and homogenous coating could be achieved (**Figure S4**), which shows that the approach is also possible for chirality pure samples. We continued with the standard (6,5)-SWCNT enriched SWCNT samples because they are more accessible to the broader community.

The samples with different densities of (GT)_10_-SWCNT coatings were tested for dopamine sensitivity by recording the time course of fluorescence intensity. These samples with varying SWCNT concentrations (0.2 nM, 1 nM, and 5 nM) exhibited different intensity increases (**Figure 2 c**) upon addition of dopamine (100 μM). This suggests that the interaction between the dopamine molecules and the number of SWCNT sensors/binding sites on the SWCNTs plays a crucial role in maximizing fluorescence change.

The test series was extended to substrates with different SWCNT concentrations. Then the maximum intensity change after dopamine addition was plotted against the starting intensity I_0_ of the SWCNT coating in PBS (**Figure 2 d**), which showed a maximum response at medium (starting) intensities. The initial rise in signal change observed with increasing SWCNT concentration can be attributed to the increased likelihood of interactions between dopamine and the sensors. However, as the absolute SWCNT concentration continues to rise, excess binding sites accumulate. As a result, a larger number of binding sites remain unoccupied, combined with an increased likelihood of quenching/reabsorption due to the higher SWCNT density. This leads to an overall decrease in absolute signal change beyond a certain SWCNT concentration. Based on this picture, a model was developed, assuming a linear increase in the number of binding sites of the SWCNT sensors with the starting intensity I_0_ of the sensors:

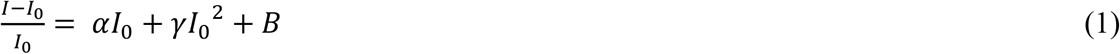

Here, the signal change is first of all proportional to the SWCNT density (I_0_) due to the rising fraction of occupied binding sites relative to the total number of binding sites. Beyond a turning point, the signal change decreases quadratically by γ*I*_0_^2^(with γ < 0), representing the fraction responsible for the density-dependent quenching/reabsorption and the oversupply of free binding sites with increasing SWCNT concentration. B accounts for background signals. Despite the complexity of the underlying processes, this model fits the experimental data in **Figure 2 d** (α = 8.95E-4, with B = 0.20 and γ = -2.57E-6 (R^2^ > 0.95)).

The most sensitive coating for dopamine detection, considering an upper concentration limit of 100 μM, can be achieved at a starting intensity I_0_ of 175 a.u. with a maximum signal change of 46%, while the smallest signal change was 4% (11.5-times lower) at I_0_ of 975 a.u. This illustrates the importance of optimizing the coating to maximize sensitivity. Since intensity values depend on the microscope and microscope settings I_0_ was additionally plotted as a function of the SWCNT concentration used for spin coating which serves as a calibration curve for the desired fluorescence starting concentration (**Figure S5**). The most sensitive coating was achieved at a concentration of 4.2 nM. The low standard deviations of I_0_ as a function of the spin-coated SWCNT concentrations indicate a highly reproducible SWCNT coating, even when comparing multiple coated samples (3 samples measured at different positions, resulting in at least n=30 positions in total). Note that manual coating of SWCNTs with a 5 nM SWCNT solution without spin coating (indicated by an asterisk in **Figure S5**) resulted in both higher intensities I_0_ and larger intensity variations. This suggests that spin coating leads to more consistent and reproducible results. It is also noticeable that the relationship between the initial intensity of the sensor coating I_0_ and the SWCNT concentration does not increase linearly, but more sharply with increasing SWCNT concentration. This trend is interesting because normally a non-linear decrease in the fluorescence signal with increasing SWCNT concentration is observed due to reabsorption effects.^[41]^ However, perhaps this effect is not as strong because the SWCNTs are immobilized and not in solution, which tends to result in greater spacing between individual SWCNTs. One possible explanation could be the preferential immobilization of long SWCNTs during the spin coating process. Due to the higher amount of long SWCNTs at higher SWCNT concentrations, longer SWCNTs may have a higher probability of being immobilized on the substrate. As longer SWCNTs have higher quantum yields compared to shorter SWCNTs^[42]^, this could account for the observed non-linear increase in SWCNT fluorescence with increasing SWCNT concentration. Another possibility is that at high concentrations physisorption of SWCNTs becomes faster, which could have an impact on the photophysical response.

### 2.2. Sensor material robustness

To ensure widespread use of a material such as Smart Slides, robustness under realistic experimental conditions has to be ensured. So far, such sensors have been used for experiments directly after preparation^[17,30,38]^ and it was unknown whether they remain functional after drying. For this purpose, sensitivity was evaluated after storing the sensor coatings in dark conditions at room temperature and in the refrigerator (**Figure 3 a**). When stored at room temperature, the sensors retained their functionality. However, after a period of 10 weeks, the intensity change observed after dopamine addition was twice as high as the immediate post-production values. If an accurate concentration determination is required when detecting a substance, this would require recalibration of the sensors to account for the increased intensity change. In contrast, the results for storage at 4°C (refrigerator) showed high stability. After one week of storage, the sensors exhibited a slight signal increase upon dopamine addition, but the intensity change remained stable for storage durations exceeding two months. It should be noted that the responses for different storage conditions could be partly due to the variance in addition of dopamine onto the Smart Slides and mixing/diffusion differences of individual experiments after a non-infinite time period. One possible reason for the altered intensity change during storage could be conformational changes in the single-stranded DNA (ssDNA) around the SWCNTs or exposure to oxygen.^[43]^

**Figure 3.**
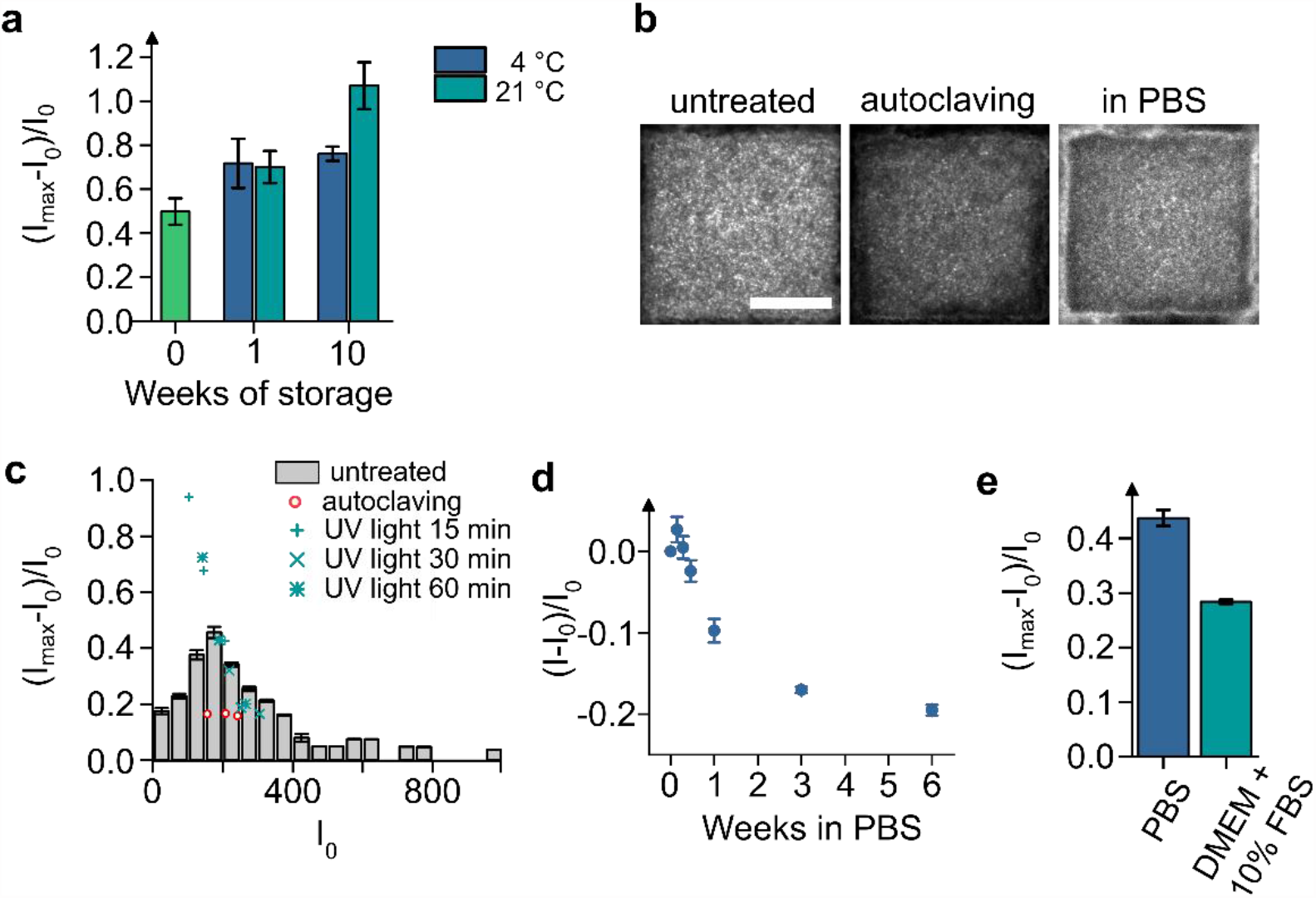
Robustness of (GT)_10_-SWCNT-based sensor coatings. a) Long-term capability to detect dopamine (100 μM) of Smart Slides stored for several weeks in the refrigerator or at room temperature (mean ± SE, n = 3). b) NIR fluorescence images before (untreated) and after autoclaving (121°C, 20 min) and in PBS (contrast adjusted as significantly darker). Scale bar = 20 μm. c) Individual sensor responses to dopamine (100 μM) after sterilization by autoclaving (red circles) and UV light (applied for 15, 30, or 60 min, blue crosses) compared to the averaged sensor responses of untreated sensor coatings (gray bars, results from Figure 2 d, mean ± SE, n ≥ 8) as a function of the initial SWCNT fluorescence intensity I_0_ d) Change of ((GT)_10_-SWCNT coating) responses over the course of multiple weeks in PBS. e) Functionality to detect dopamine (100 μM) in full cell culture medium on a Smart Slide compared to PBS. (mean ± SE, n = 3).

The more stable results observed at 4°C storage align with general recommendations for storing ssDNA because refrigeration increases the stability of ssDNA by at least twice the time compared to storage at room temperature.^[44]^

To assess the functionality of the sensors after typical sterilization procedures autoclaving and UV light were used (**Figure 3 b, c**). Autoclaving of the sensors involved a preliminary experiment to determine whether the SWCNT sensors in dispersion are resistant to heating. (GT)_10_-SWCNTs, (GT)_40_-SWCNTs, and locked nucleic acid (LNA) (GT)_15_-SWCNTs are known for their lower flexibility and potentially higher thermal stability. They were subjected to heating at 60°C, 80°C, and 100°C for 30 minutes to test if they are stable at these temperatures without aggregation (**Figure S6 a**). Both, (GT)_10_- and LNA (GT)_15_-SWCNTs precipitated when heated to 100°C, while (GT)_40_-SWCNTs showed increased stability and remained in solution. This is consistent with observations that SWCNTs with longer ssDNA sequences have higher thermal stability.^[45]^ In contrast, dry (GT)_10_-SWCNT coatings heated to 100°C did not exhibit any limitations in functionality and responded with a fluorescence increase upon the addition of 100 μM dopamine (**Figure S6 b**). Thus, sensor coatings demonstrated superior stability compared to SWCNT dispersions, which suggests that autoclaving as a sterilization method could be possible.

To assess the adhesion of the sensor coating during autoclaving, the sensors were spin-coated onto glass substrates with an imprinted location grid. However, the grid did not allow homogeneous coating as on smooth glass substrates (**Figure S7 a**). After autoclaving, slight changes in the coating appeared on the surface. In some cases, there were fewer sensors and the surface appeared darker, while in others it appeared brighter. The addition of PBS quenched the fluorescence of the sensors, but also led to the detachment of SWCNTs, resulting in weaker responses to dopamine (**Figure S7 c**). To improve the adhesion of SWCNTs, an additional baking step (heating the silanized substrates at 120°C for 2 h) was introduced after the aminosilane and before the SWCNT coating process to remove physisorbed silane molecules, which does not affect covalently bonded silane molecules.^[46]^ The SWCNT coating on the treated grids demonstrated significant improvements in homogeneity, adhesion after autoclaving, and sensitivity after dopamine addition (**Figure 3 b, Figure S7 b, c**). Without the additional heating step of the APTES layer, the non-covalently bound APTES molecules most likely desorbed during autoclaving, and thus most SWCNTs detached after the addition of PBS. A comparison of the starting intensities (**Figure 3 c**) revealed that sensitivity was reduced by approximately 50% compared to the non-autoclaved grids. Covalent functionalization methods^[47]^ and/or immobilization via avidin-biotin interaction^[36]^ could further improve the stability of the system.

Sterilization of the sensor coatings using UV light was performed with different exposure times (15, 30, and 60 minutes), followed by testing their functionality to detect dopamine. In most tests, the functionality of the sensors remained unaffected after UV exposure, regardless of the exposure duration, when compared to the untreated samples (**Figure 3 c**). Only sensors with lower starting intensity I_0_ showed larger signal changes than the untreated samples, although the reason for this observation remains unclear. The absorption of UV light can lead to the formation of pyrimidine dimers, primarily involving adjacent T-T and T-C sequences.^[43]^ C-T and C-C sequences are less prone to photoreactivity. In rare cases, UV radiation can induce modifications in DNA purine bases, including adenine residues undergoing photocycloaddition reactions with adjacent adenine or thymine. The (GT)_10_ sequences used in the sensor coatings are expected to be largely unaffected by UV radiation. It is possible that other DNA sequences could be more susceptible to the effects of UV radiation, as dimer formation can lead to changes in DNA conformation and a loss of sensitivity. However, in the specific coating region that was determined to be the most sensitive, the sensor responses remained unchanged for different exposure times. As a result, a 30-minute UV exposure was chosen for subsequent cell experiments as sterilization method. We refrained from testing other used sterilization methods such as gamma radiation and ethylene oxide because glass discoloration occurs when sterilizing with gamma radiation^[48]^ and autoclaving and UV irradiation are cheaper, faster and easier sterilization options compared to ethylene oxide. In addition, there are uncertainties regarding the risk of toxicity from ethylene oxide residues.^[49,50]^

Next, long-term adhesion of the sensors in liquid was studied, which is important for cell experiments requiring cell cultivation over several weeks. Signals decreased by approximately 20% after 6 weeks (**Figure 4 d**). Adhesion was compared in PBS and water, as well as in PBS after the aforementioned aminosilane coating with an additional baking step (**Figure S8**). Contrary to expectations, the last-mentioned variant exhibited lower long-term adhesion, with hardly any SWCNTs remaining after only 3 weeks. As mentioned earlier, the use of avidin-biotin interactions or covalent immobilization strategies could potentially enhance the stability of the sensors in liquid and improve long-term stability. However, for most cell experiments less than one week of cultivation is sufficient and the observed reduction in signal after 6 weeks should not pose a significant issue.

**Figure 4.**
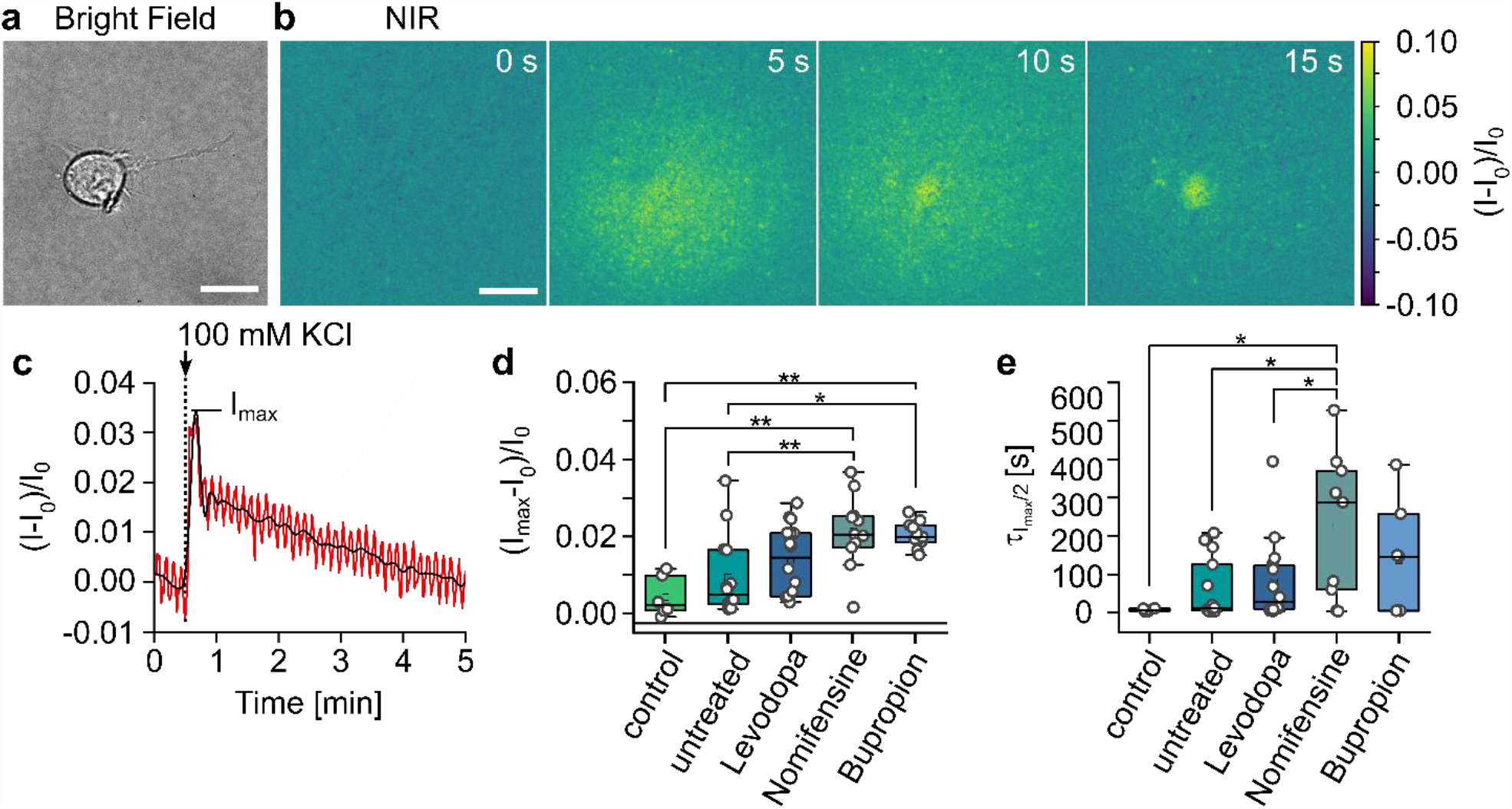
Smart Slides to monitor cell responses to drugs. a) Bright field image of a Neuro 2a cell cultured on a (GT)_10_-SWCNTs coated glass surface. b) Color-coded NIR intensity changes of (GT)_10_-SWCNTs at different time points after stimulation of dopamine release with 100 mM KCl. All scale bars = 20 μm. c) Typical intensity change of the sensor layer under a single cell. D) Maximum intensity changes and e) decay time of the maximum intensity changes on I_max_/2 of SWCNTs after KCl stimulation of controls (without cells), untreated cells, and cells treated with different drugs (5 μM for 10 min, n=6 for control, n≥11 cells for the rest). Note that not for all experiments the value 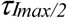 could be determined from the time traces, so that n=5 for control, n≥7 cells for the rest. Statistically significant differences marked with * p≤0.05, ** p≤0.01, *** p≤0.001 (ANOVA and Tukey test). Not significant differences are not indicated.

Finally, the sensitivity of the sensors was evaluated in cell medium supplemented with 10% fetal bovine serum (FBS). Detection was possible in cell medium but sensitivity was reduced by approximately one-third compared to the simple buffer environment (**Figure 3 e**). The reduced sensitivity in cell medium can be attributed to additional compounds in the cell medium that interact with the sensors and potentially interfere with the binding of dopamine, leading to a diminished fluorescence response. To improve the sensitivity of the sensors in protein-rich environments, a detailed understanding of the interaction between the nanosensors and their environment is required.^[51]^ In addition, covalent linking the SWCNTs to the surface could increase robustness by reducing desorption.^[22]^ Alternatively, for the cell experiments, the cell medium could be replaced by simpler (low-serum) buffers.

### 2.3. Monitoring of Biochemical Cell Responses

Smart Slides should be able to map dopaminergic release processes of neuronal cells, providing access to a better understanding of cellular communication. Furthermore, they have potential applications in drug development, allowing the measurement of the effects of different substances on dopamine release in the synaptic cleft. Here, differentiated Neuro-2a cells, a standard dopaminergic cell model for neurodegenerative diseases, were directly seeded onto the sensors. After approximately 18 h, the cells exhibited good adhesion without additional coating (**Figure S9**). In order to evaluate the effectiveness of the Smart Slides, different dopaminergic substances, including levodopa (commonly used to increase dopamine levels in Parkinson’s disease patients), nomifensine, and bupropion (both dopamine-norepinephrine reuptake inhibitors in antidepressant therapy), were added to the cell culture for 10 min. Since it is known that such sensors also respond to dopamine homologues^[25]^, it was tested whether (GT)_10_-SWCNTs also elicit a sensor response by the added substances in solution (**Figure S10**). Levodopa affects the fluorescence of (GT)_10_-SWCNTs due to its chemically similar structure. However, the response is only half as strong compared to dopamine, while nomifensine and bupropion elicit slightly negative sensor responses. To ensure that the sensors were not biased during dopamine measurements, cells were washed twice with PBS before a third time PBS supplemented with calcium chloride was added to start the release experiments. Subsequently, spatial and temporal dopamine release from the cells was measured by stimulating exocytosis with potassium chloride (KCl).

**Figure 4 a - c** illustrates the results of an untreated cell experiment where no substances were added. The bright field image shows a cell adhering on the sensor coating, while the color-coded image represents the NIR channel before and after the stimulus was added. Immediately after the stimulus, there is a localized increase in fluorescence intensity around the cell, which subsequently diminishes due to the diffusion of released dopamine. For better comparison, the cell area was selected in each experiment to calculate the mean intensity changes over time (**Figure 4 c**). Analysis of all individual experiments revealed some heterogeneity, wherein not all cells demonstrated dopamine release. This variability in response is consistent with other studies measuring cellular dopamine release.^[31,52,53]^ Moreover, the sensors exhibited expected differences in signal change following treatment with dopaminergic substances. Selected measurements for the different conditions are provided for comparison in **Figure S11**. For instance, nomifensine and bupropion tended to show amplified and prolonged signal changes following stimulation, which can be attributed to their role as dopamine reuptake inhibitors, prolonging the presence of dopamine in the synaptic cleft.

To account for differences in cellular responses across experiments, the maximum intensity change I_max_ and the time required for the signal change to decrease from I_max_ to I_max_/2 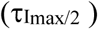 were extracted and presented as a box plot based on the respective cell treatment conditions (**Figure 4 d, e**). Additionally, a box blot was created to depict the time difference between stimulation and maximum signal change I_max_ (**Figure S12**).

Note that the time- and spatially-resolved images and time traces contain even more information that could further be extracted. Additionally, the rate constants of the sensors are important to further interpret the kinetics of the chemical images, as the temporally and spatially resolved images are influenced by the rate constants/kinetics of the sensors^[54]^. The population mean changes were found to be significantly different. Control experiments for which KCl was added to the Smart Slides without cultured cells resulted in minimal or no fluorescence changes, which can be attributed to the local increase in ion concentration.^[55]^ To address this issue, LNA nanosensors can increase stability to ion-induced perturbations^[56]^ or optogenetic stimulation can be used for more specific and controlled stimulation.

Addition of levodopa to the cells resulted in an increase in dopamine release, as evidenced by a 193% increase in the median maximum signal change and a 152% increase in 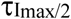 compared to untreated cells, indicating higher dopamine concentrations. The overall increase is consistent with previous studies, although the absolute values differ, which can be attributed to differences in the cellular system, concentration, and exposure time of levodopa.^[52,57,58]^ The addition of nomifensine and bupropion increased the median maximum signal by 300%, with 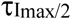 increasing from 11 s for untreated cells to longer-lasting presence times of dopamine in the synaptic cleft (288 s for nomifensine and 146 s for bupropion).

For bupropion, the extracted values of 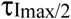 are not significant compared to untreated cells, which is probably due to the fact that not all bupropion (and partially also nomifensine) cell experiments could be used to evaluate I_max_/2, as the signal increase did not drop half of I_max_ within the 10-minute experimental duration. In some cases, the maximum signal change was measured several minutes after stimulation (**Figure S12**). Amperometric measurements in bovine chromaffin cells treated with 5 μM bupropion also showed significant increases in I_max_ level (around 151%).^[59]^ However, no influence on the release kinetics was observed, as the half-width of an amperometric spike remained unchanged^[59]^, while Nomifensine-treated PC12 cells exhibited prolonged times of dopamine presence in the synaptic cleft.^[60]^

Overall, these findings demonstrate the effectiveness of using Smart Slides to monitor cellular release processes and their modulation by the addition of neuropharmaceutical drugs. This approach can easily be extended to other targets/biological questions, such as coagulation disorders^[61]^, diabetes^[62,63]^, cancer^[64]^, hydrolytic enzyme activity^[65]^, or inflammatory markers.^[66]^

## 3. Conclusion

In summary, we tailored NIR fluorescent SWCNTs for dopamine detection and optimized the coating of these sensors on surfaces, resulting in functional and highly sensitive Smart Slides. These slides showed the necessary robustness in terms of their long-term stability during storage and in liquids for several weeks, as well as their compatibility with UV light sterilization. Furthermore, we demonstrated their applicability in drug testing by directly measuring the effects of dopaminergic compounds on dopamine release. This approach is not limited to the detection of dopamine but can be extended to other targets and biological questions due to the easily customizable surface chemistry of the SWCNTs and the wide range of SWCNT sensors already developed. Overall, the development of these Smart Slides represents a significant advance in the field of cellular analysis by providing a simple tool that can access optically dynamic information about cellular communication and drug effects in a controlled and standardized manner.

## 4. Experimental Section/Methods

### Materials

All materials were purchased from *Sigma-Aldrich* unless specified otherwise.

### SWCNT Surface Modification

For surface modification of (6,5) chirality-enriched CoMoCAT-SWCNTs (Sigma-Aldrich, product no. 773 735) with ssDNA, a recently published protocol was used. ^[55]^ SWCNTs (100 μL, 2 mg mL^−1^ in PBS (Carl Roth)) were mixed with ssDNA (100 μL, 2 mg mL^−1^ in PBS), followed by tip sonication (ice bath, 10 min at 30% amplitude, Fisherbrand Model 120 Sonic Dismembrator) and centrifugation (2 × 30 min, 16100 g, 4°C, Hettich MIKRO 200R) to remove aggregates. The supernatant yielded homogenously dispersed (GT)_10_-SWCNTs, which were diluted in PBS to the desired SWCNT concentration after determination of SWCNT concentration *via* absorption measurements (see NIR Spectroscopy).

### NIR Spectroscopy

Absorption spectra for the characterization of SWCNT samples were recorded using a JASCO V-780-ST spectrophotometer in the wavelength range of 400– 1350 nm in 0.5 nm steps in quartz cuvettes (Hellma, 10 mm optical path). SWCNT concentration was calculated based on previous literature.^[55,67–69]^

Fluorescence spectra in the range between 835 – 1245 nm were recorded with 5 s integration time using a spectrometer (Shamrock 193i, Andor Technology Ltd.) connected to a microscope (IX73, Olympus) equipped with a 20x objective (plan achromat infrared objective LCPLN20XIR) and a 561 nm laser at 100 mW (Gem561, Laser Quantum) for excitation.

### Raman Spectroscopy

Raman measurements were carried out using a confocal Raman microscope (inVia InSpect from Renishaw) equipped with a 100x objective and an excitation of 532 nm (25 mW laser power) with an integration time of 10 s.

### SWCNT-coated surfaces

35mm Glass bottom petri dishes (ibidi) were treated with oxygen plasma (Atto B, Diener electronic, 0.6 mbar, 1 min). Directly after plasma cleaning, surfaces were coated with APTES solution (300 μL of 1 wt% APTES/H_2_O in ethanol) and incubated for 1 h at room temperature. Subsequently, petri dishes were washed with ethanol, rinsed with H_2_O, and then either dried directly with N_2_ or baked at 120°C for 2 h, followed by rinsing again with H_2_O and drying with N_2_. SWCNT coating was performed by spin coating (Laurell WS-650MZ-8NPP) at 1600 rpm for 60 s.135 μl of diluted (GT)_10_-SWCNT solution was added directly after starting the rotation. Parameter selection was guided by Card et al.^[70]^ To remove non-immobilized SWCNTs, the surface was rinsed with H_2_O and dried with N_2_.

### Imaging of Single SWCNT sensor coatings

For characterization of SWCNT-coated surfaces and dopamine response measurements an inverted microscope (Nikon Eclipse Ti2) equipped with a 100× oil immersion objective (CFI Plan Apochromat Lambda D 100x Oil/1.45/0.13) was used. A white LED (CoolLED pE300 Lite, 100% power) was used in combination with a 560 ± 40 nm bandpass filter (AHF analysentechnik F47-561) for the excitation of SWCNTs via the E_22_ transition. Excitation light was eliminated from emission via a dichroic mirror (transmission > 93% at 813.5 nm, AHF analysentechnik F38-801) and an 840 nm long-pass filter (AHF analysentechnik F47-841). Images were taken with a Si camera (Hamamatsu Orca Flash 4.0) with 2048 × 2048 pixels at 1 s integration time.

For dopamine response measurements, 20 μL of a freshly prepared dopamine solution (dopamine hydrochloride in 1× PBS) was added to 2 mL of PBS/cell medium in the petri dish after a baseline I_0_ was recorded for 30 s. The final dopamine concentration was 100 μM. For characterization of the resulting intensity change (I−I_0_)/I_0_ after dopamine addition, the initial (I_0_) and final intensity after dopamine addition (I) were calculated by averaging the respective frames of the recorded video. Note that the SWCNT coating optimization was performed within this standard microscope with a LED and a Si camera for excitation and detection. In the later cell experiments, which were performed in a different setup with a laser and a NIR camera (see Cell Experiments), very low laser powers in the range of 55 and 120 mW were required, which means that the detection should also be possible with “standard” equipment using (6,4)-(GT)_10_-SWCNTs for gaining even higher sensitivity.^[33]^

### Robustness tests

Storage of (GT)_10_-SWCNT-coated petri dishes was performed after drying the dishes with N_2_ either at room temperature in a dark box or a refrigerator at 4°C. To test sterilization capabilities, SWCNT-coated petri dishes were either autoclaved at 121°C and 100% humidity (2 bar) for 20 min (Espec EHS-211 M) or irradiated with 254 nm UV light in a biosafety cabinet (HMC Europe BSC-700IIA2-G) for 15, 30 or 60 min. To test the stability of the SWCNT coating in liquid, SWCNTs were coated on an imaging dish with a glass bottom and an imprinted 50 μm grid (ibidi) to ensure easy imaging of the same SWCNT coated location on different days. Petri dishes were filled with 2 mL of the respective liquid. A lid and sealing with parafilm ensured that evaporation of the liquid was minimized.

### Cell Culturing

NEURO-2A cells were purchased from DSMZ German Collection of Microorganisms and Cell Cultures (ACC 148) and cultivated according to the supplier’s protocol in a humidified 5% CO_2_ atmosphere at 37°C in T-75 flasks (Sarstedt) with a sub-cultivation ratio of 1:4 every 5-7 days. Cells were grown in 16 mL DMEM (Thermo Fisher Scientific) supplemented with fetal bovine serum (FBS) (10%), 1x non-essential amino acids, penicillin (100 units mL^−1^), and streptomycin (100 μg mL^−1^, Thermo Fisher Scientific). Differentiation of NEURO-2A cells was performed according to Tremblay *et al*.^*[71]*^ Prior to experiments (96 hours), cells were cultivated in DMEM supplemented with 1 mM dibutyryl cyclic adenosine monophosphate (dbcAMP) and 0.5% FBS. After three days cells were carefully rinsed with medium from the surface of cultivation flasks and a total number of 75,000 cells was transferred to SWCNT-coated glass surfaces, which had previously been sterilized with 30 min UV exposure, and allowed to adhere for 18 hours.

### Cell Experiments

The setup used to perform the cell experiments consisted of an inverted microscope (Olympus IX73) equipped with a 100x oil immersion objective (UPlanSApo/1.35/0.13-0.19). A 561 nm (Jive 500, Cobolt) at 55 or 120 mW was used for SWCNT excitation. Excitation light was eliminated from emission via a 900 nm long pass filter (Thorlabs FELH0900). The NIR fluorescence was imaged with a NIR camera (Xenics Cheetah-640-TE1 InGaAs camera) at 1 s exposure time. Bright field images of cells were taken with a Si camera (pco.panda).

For dopamine release experiments, the cell medium was exchanged for PBS (1 mL) supplemented with MgCl_2_ and CaCl_2_. To stimulate dopamine release, a KCl solution (34 μL of 3 M) was added after recording a 30 s NIR fluorescence baseline, resulting in a final concentration of 100 mM KCl. Cells and PBS (Carl Roth) were kept at 37°C before the experiment, assuming a temperature close to 37°C during the cell experiments.

To assess the impact of dopaminergic substances on dopamine release, 10 min before the start of the cell experiment, the specific substance of interest (Levodopa, Nomifensine maleate salt (Medchem Express, Fisher Scientific), Bupropion hydrochloride (Enzo Life Sciences, Fisher Scientific) diluted in PBS was added to the cell medium (20 μL to 600 μL cell medium), resulting in a concentration of 5 μM. After 10 min of exposure, the cell medium was carefully aspirated and the cells were rinsed twice with PBS. Subsequently, 1 mL of PBS was added to perform the imaging and stimulation with KCl. In control experiments, 100 mM KCl was added to the Smart Slides containing 1 mL PBS without cultured cells.

### Data Processing and statistical analysis

Fiji was used for image data processing and OriginPro for statistical analysis. To determine the intensity changes of SWCNT fluorescence in the cell experiments, the average intensity of the cell area was selected for analysis in each case. The first 30 frames, which served to record the baseline I_0_, were averaged and the resulting time-dependent signal change (I-I_0_)/I_0_ was calculated. The signal change was smoothed (OriginPro, Wavelet DB6). From these spectra, the maximum signal change Imax and the decay time σ_Imax/2_ were extracted. A one-way analysis of variance (ANOVA) was conducted to examine statistically significant differences among the various treated cell groups, using a significance level of 0.05. Post-hoc comparisons of mean values were performed using the Tukey test.

## Acknowledgements

This work was supported by the Fraunhofer Internal Programs under Grant No. Attract 038– 610097. Funded by the Deutsche Forschungsgemeinschaft (DFG, German Research Foundation) under Germany’s Excellence Strategy – EXC 2033 – 390677874 – RESOLV. This work was supported by the “Center for Solvation Science ZEMOS” funded by the German Federal Ministry of Education and Research BMBF and by the Ministry of Culture and Research of Nord Rhine-Westphalia. This work was funded by the VW foundation.

