## Supplemental Figures for "Smart Slides for Optical Monitoring of Cellular Processes"

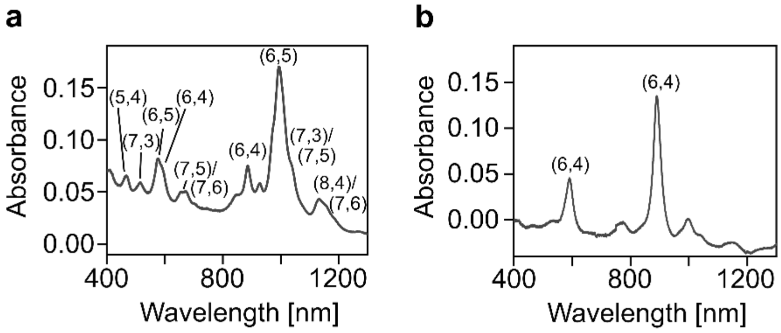


***Figure S1. Absorbance spectra of SWCNTs.*** *a) (GT)$_{10}$-CoMoCAT-SWCNTs, b) (GT)$_{10}$-(6,4)-SWCNTs with assigned chiralities.*

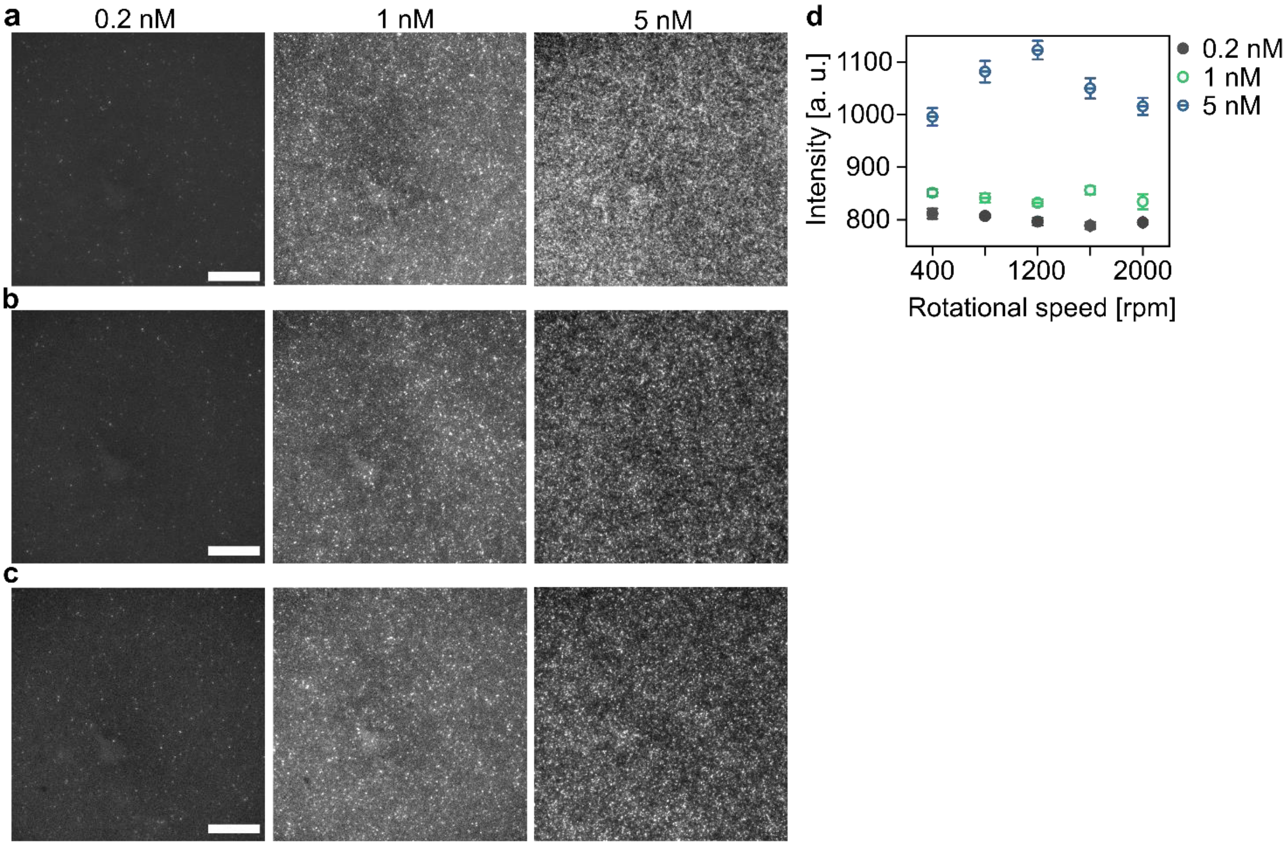


***Figure S2. NIR fluorescence images of (GT)$_{10}$-SWCNT coatings.*** *Prepared by spin coating at a) 400 rpm, b) 1200 rpm and c) 2000 rpm with different SWCNT suspension concentrations. Scale bars = 20 µm. d) SWCNT fluorescence intensity of (GT)$_{10}$-SWCNT coatings of different concentrations as a function of the rotational speed used for spin coating (mean ± SE, n=28).*

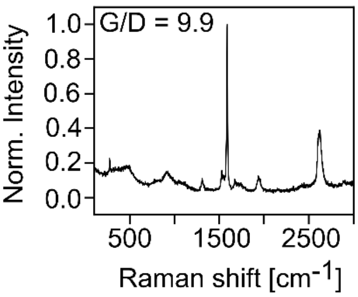


***Figure S3. Raman spectrum of a (GT)$_{10}$-SWCNT coated surface.*** *The G/D ratio indicates a low defect level in the SWCNTs.*

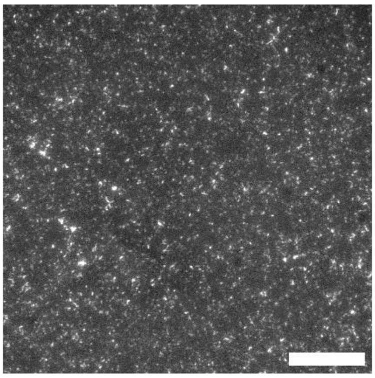

***Figure S4. NIR image of spin coated (GT)$_{10}$-(6,4)-SWCNTs.*** *135 µl of a 0.1 nM SWCNT solution was spin coated six times in succession at 1600 rpm to obtain a dense coating. Scale bar = 20 µm.*

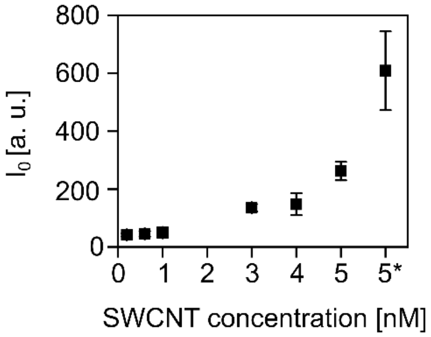


***Figure S5. Intensity calibration curve.*** *Initial SWCNT fluorescence intensity $I_0$ of (GT)$_{10}$-SWCNT coatings in PBS as a function of the SWCNT concentration used for spin coating. Note 5* represents manual coating of 5 nM SWCNTs without spin coating (mean ± SD, n>30).*

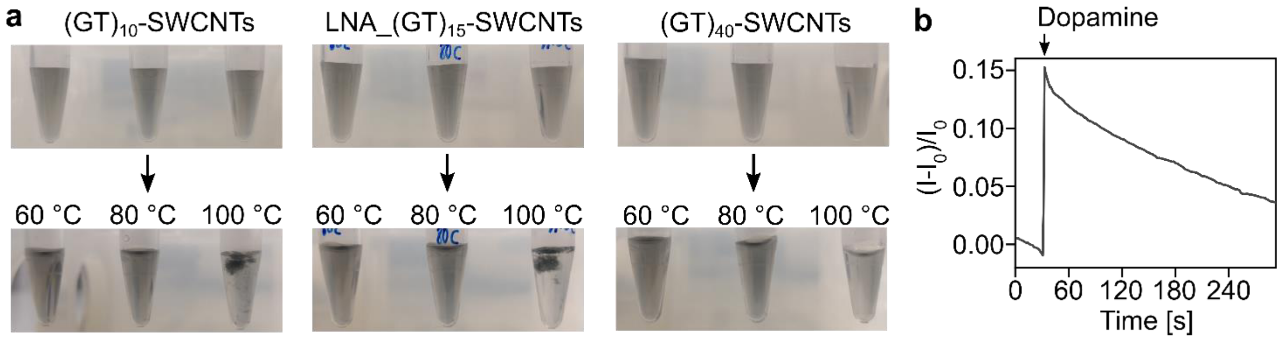


***Figure S6. Heating of ssDNA-SWCNTs.*** *a) Images of different ssDNA-SWCNT suspensions ((GT)$_{10}$-, locked nucleic acid (LNA) (GT)$_{15}$- and (GT)$_{40}$-SWCNTs) before and after heating (60°C, 80°C, 100°C for 30 min). With the exception of (GT)$_{40}$-SWCNTs, ssDNA-SWCNTs precipitate when heated to 100°C. b) Immobilized (GT)$_{10}$-SWCNTs remain functional to detect 100 µM dopamine after heating to 100°C for 30 min.*

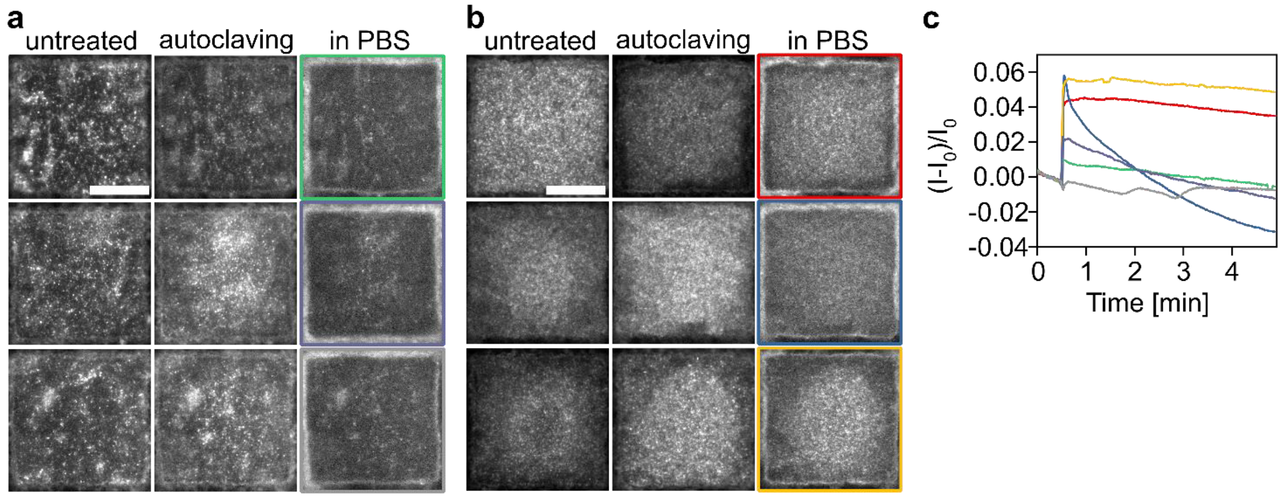


***Figure S7. Autoclaving of immobilized ssDNA-SWCNTs.*** *Images of immobilized (GT)$_{10}$-SWCNTs before (untreated) and after autoclaving (121°C, 20 min) and in PBS. APTES coated surface a) rinsed with ethanol and water, b) additionally heated for 2 h at 120 °C before SWCNTs coating. c) 100 µM dopamine detection with these SWCNT coatings (assignment via respective colors). Scale bars = 20 µm.*

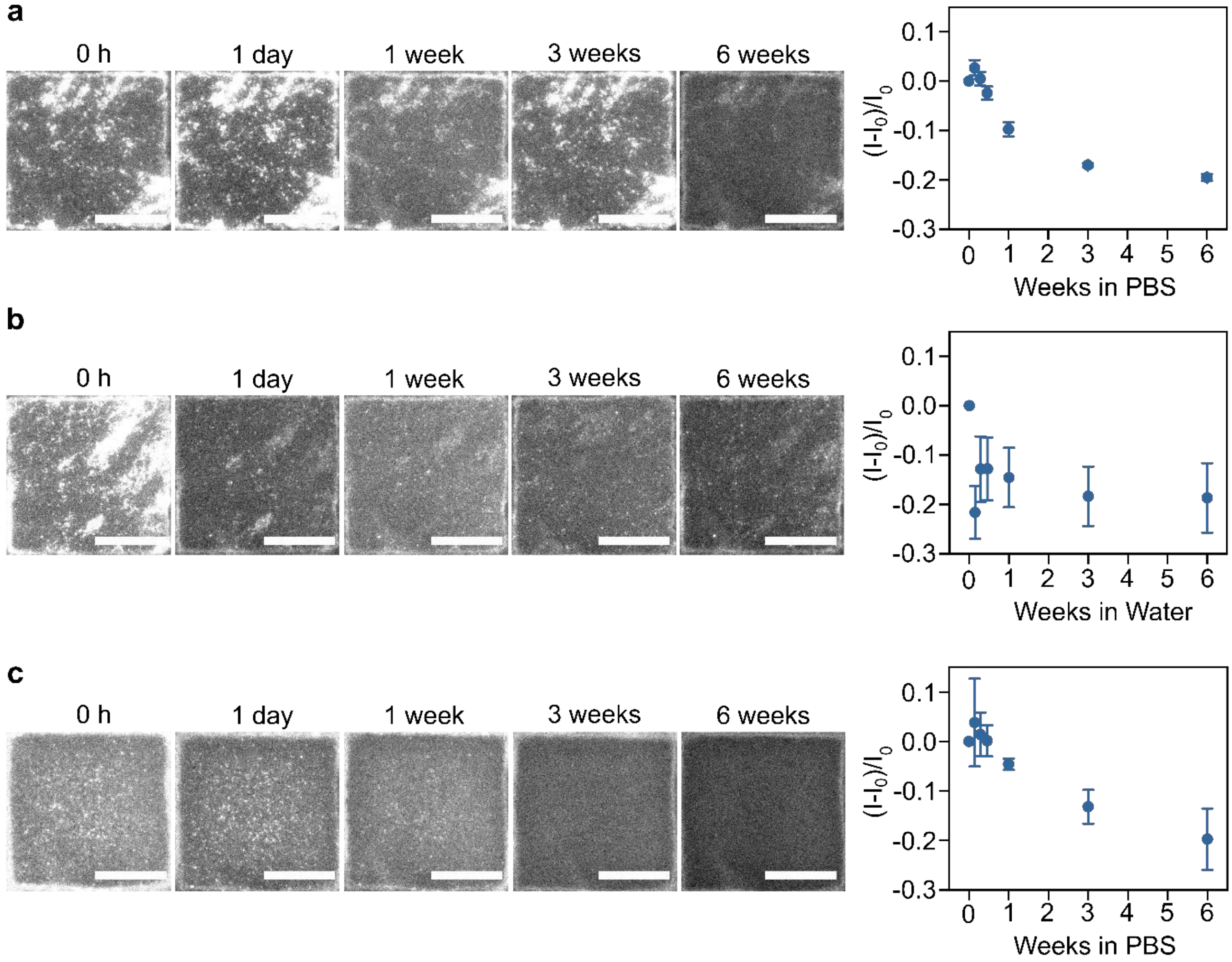


***Figure S8. Stability of immobilized ssDNA-SWCNTs in liquid.*** *NIR Images and intensity changes of the same position on 1% APTES-treated glass surfaces at different time points. a) In PBS, b) in water, c) in PBS with APTES-treated glass surfaces preheated for 2 h at 120°C (mean ± SE, n = 3). All scale bars = 20 µm.*

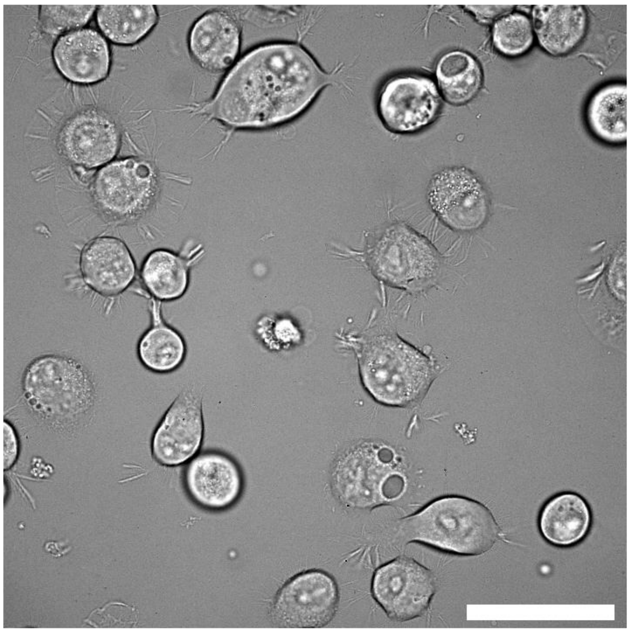

***Figure S9. Bright field image of differentiated Neuro2a cells adhering to the $(GT)_{10}$-SWCNT sensor coating.*** *Scale bar = 50 µm.*

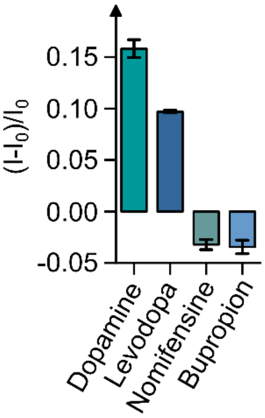


***Figure S10. Sensor response of $(GT)_{10}$-SWCNTs in solution.*** *After addition of 5 µM dopamine in comparison to the tested dopaminergic substances (levodopa, nomifensine, and bupropion, mean ± SE, n=3).*

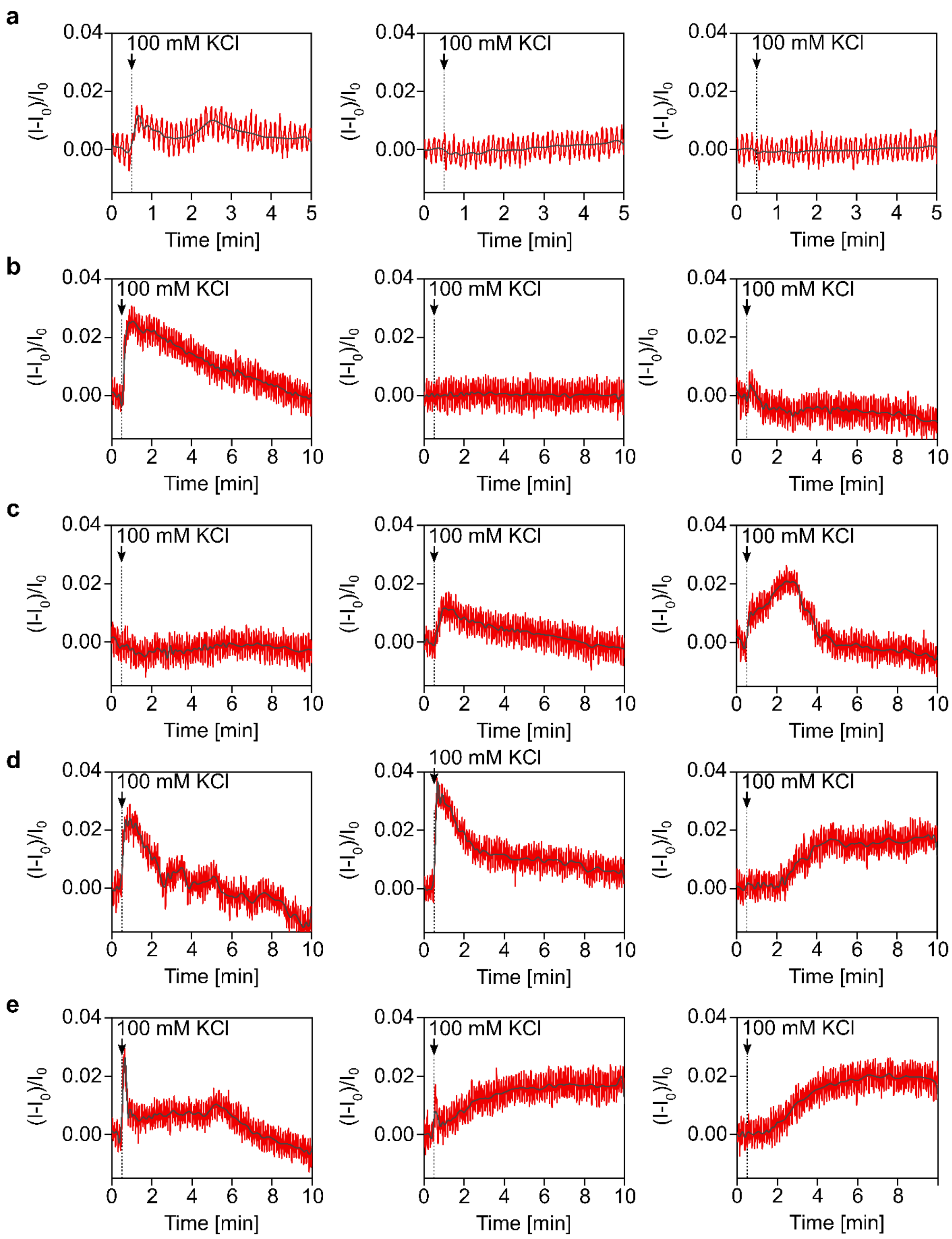


***Figure S11. Mean SWCNT intensity change of different experiments.*** *During addition of 100 mM KCl a) without cells (control), b) with untreated cells and c) after addition of 5 µM levodopa, d) nomifensine, and e) bupropion.*

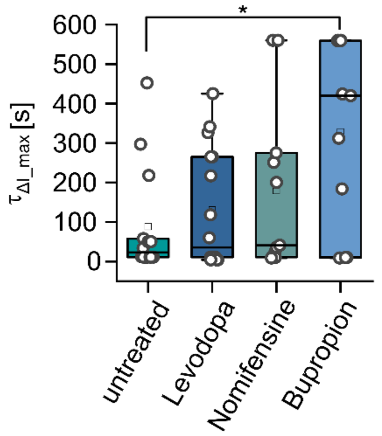


***Figure S12.*** *Box plot of the time difference between stimulation and maximum signal change $I_{max}$ of untreated cells and cells treated with different drugs (5 µM for 10 min, n=5 for control, n≥11 cells for the rest. Statistically significant differences marked with * p≤0.05.*
